# *WUSCHEL*-Related Homeobox Genes Cooperate with Cytokinin Signalling to Promote Bulbil Formation in *Lilium lancifolium*

**DOI:** 10.1101/2021.10.07.463561

**Authors:** Guoren He, Yuwei Cao, Jing Wang, Meng Song, Mengmeng Bi, Yuchao Tang, Leifeng Xu, Panpan Yang, Jun Ming

## Abstract

The bulbil is an important vegetative reproductive organ in triploid *Lilium lancifolium*. Based on our previously obtained transcriptome data, we screened two *WUSCHCEL-related homeobox* (*WOX*) genes closely related to bulbil formation, *LlWOX9* and *LlWOX11*. However, the biological functions and regulatory mechanisms of *LlWOX9* and *LlWOX11* are unclear. In this study, we cloned the full-length coding sequences of *LlWOX9* and *LlWOX11*. Transgenic *Arabidopsis* showed increased branch numbers, and the overexpression of *LlWOX9* and *LlWOX11* in stem segments promoted bulbil formation, while the silencing of *LlWOX9* and *LlWOX11* inhibited bulbil formation, indicating that *LlWOX9* and *LlWOX11* are positive regulators of bulbil formation. Cytokinins acting through type-B response regulators (type-B RRs) could bind to the promoters of *LlWOX9* and *LlWOX11* and promote their transcription. LlWOX11 could enhance cytokinin pathway signalling by inhibiting the transcription of type-A *LlRR9*. Our study enriches the understanding of the regulation of plant development by the *WOX* gene family and lays a foundation for further research on the molecular mechanism of bulbil formation in lily.

## Introduction

*Lilium lancifolium*, also known as tiger lily, is an important *Lilium* species of the Liliaceae family. *L. lancifolium* shows high adaptability and is widely cultivated in China for its edible bulbs and medicinal applications (Liang & Tamura, 2000; China Pharmacopoeia Committee, 2005; Yu *et al*., 2015), with a production value of approximately six billion Yuan per year. *L. lancifolium* is a natural triploid and cannot be propagated sexually, but its leaf axils can form a large number of purple-black bulbils (Bach & Sochacki, 2012; Chung *et al*., 2015).

Bulbils grow on leaf axils and can naturally fall off the mother plant and develop into a new complete individual after maturity (Yang *et al*., 2017). The bulbil propagation strategy has the advantages of high efficiency and better retention of maternal genetic characteristics and is therefore the main reproductive strategy for *L. lancifolium*.

Bulbils are a special and important type of reproductive organ in plants and are only formed in a few plant species, such as *Dioscorea batatas*, *Allium sativum*, *Titanotrichum oldhamii*, *Pinellia ternate*, *Agave tequilana*, and *Lilium* species (Wang *et al*., 2004; Bell & Bryan, 2008; Abraham-Juárez *et al*., 2010; Sandoval *et al*., 2012; Yang *et al*., 2017). The formation of bulbils is a complex developmental process that is regulated by genetic and environmental factors and phytohormones.

Plant hormones, especially auxin and cytokinin, have been proven to be involved in the regulation of bulbil formation, in which auxin inhibits bulbil formation, whereas cytokinin promotes the bulbil formation (Wang & Cronk, 2003; Peng *et al*., 2005; Abraham-Juárez *et al*., 2015; Navarro *et al*., 2015; He *et al*., 2020). Before bulbil initiation in *D. polystachya*, auxin rapidly accumulates in the leaf axil, followed by the expression of auxin transport genes, such as *ARF9*, *ARF18*, *AX15A*, and *AUX22D*, resulting in auxin outflow from the leaf axil and bulbil initiation (Wu *et al*., 2020). *AtqPIN1* and *AtqSoPIN1* participate in auxin outflow in *A. tequilana* (Abraham-Juárez *et al*., 2015). In *D. polystachya*, the expression of cytokinin oxidase/dehydrogenase genes (*CKX1*, *CKX3*, *CKX9* and *CKX11*) is decreased before bulblet initiation and leads to the accumulation of cytokinin in the leaf axil (Wu *et al*., 2020). In a previous study, we revealed that iP-type cytokinins were the most important cytokinins during bulbil formation and showed that the accumulation of iP-type cytokinins was mainly due to the upregulation of cytokinin biosynthesis genes (*IPT1* and *IPT5*) and cytokinin activation genes (*LOG1*, *LOG3*, *LOG5* and *LOG7*) and the significant downregulation of cytokinin degradation gene (*CKX4*) expression (He *et al*., 2020).

As a special type of axillary organ, bulbils originate from the axillary meristem (AM). A recent study revealed that cytokinins can promote AM initiation through cytokinin type-B response regulators (type-B RRs) (Wang *et al*., 2017). Type-B RRs are positive regulatory transcription factors in cytokinin signalling and mostly modulate the transcription of cytokinin-regulated genes by directly binding target DNA sequences at their C-terminal MYB domains (Hosoda *et al*., 2002; Kieber & Schaller, 2014). In *A. thaliana,* cytokinin signalling is mainly mediated by five members of type-B RR subfamily I: ARR1, ARR2, ARR10, ARR11 and ARR12 (Mason *et al*., 2004, 2005; Schaller *et al*., 2007; Yokoyama *et al*., 2007; Ishida *et al*., 2008; Tsai *et al*., 2012). In regulating axillary bud formation, type-B RRs act as key transcriptional regulators involved in AM initiation. ARR1 can directly bind to the *WUS* promoter and activate the transcription of *WUS*; ARR2, ARR10, ARR11 and ARR12 can also activate the expression of *WUS*, indicating that type-B ARRs show functional redundancy in regulating the expression of *WUS*, in which ARR1 is the key regulatory factor (Wang *et al*., 2014a,b, 2017).

Regarding the molecular regulation of bulbil formation, however, only a small number of genes related to bulbil formation have been identified to date, and the associated regulatory mechanism is not clear. In *A. tequilana*, *AtqKNOX1* and *AtqKNOX2* are expressed at the beginning of globular bud formation and are specifically expressed during meristem development (Abraham-Juárez *et al*., 2010). The expression of some *AtqMADS* genes is decreased during bulbil formation, indicating that *AtqMADS* genes may be negatively related to bulbil formation in this species (Sandoval *et al*., 2012). In *T. oldhamii*, the expression of *Gesneriaceae-FLORICAULA* (*GFLO*) is also downregulated during bulbil formation, indicating that *GFLO* acts as a negative regulator during bulbil formation (Wang *et al*., 2004). The AGO protein mediates the silencing of downstream genes through miRNA. In *L. lancifolium*, *LlAGO1* is specifically expressed in the bulbil and upregulated during bulbil formation, which indicates that the miRNA pathway may also be involved in the regulation of bulbil formation (Yang *et al*., 2018).

The WUSCHEL-related homeobox (WOX) proteins are a plant-specific family within the eukaryotic homeobox transcription actor superfamily characterized by a conserved N-terminal homeodomain (HD) consisting of 60-66 amino acids (Mayer *et al*., 1998; Haecker *et al*., 2004). Functional studies have revealed that the WOX transcription factors play important roles in promoting cell division, preventing immature cells from differentiating, embryonic development, stem cell niche maintenance in the meristem and organ formation (Stahl *et al*., 2009; Van Der Graaff *et al*., 2009; Yadav *et al*., 2011). Based on the phylogenetic analysis and the distribution of *WOX* genes in the plant kingdom, they have been classified into three clades: a modern/WUS clade (found in seed plants), an intermediate/WOX9 clade (found in vascular plants including lycophytes), and an ancient/WOX13 clade (found in vascular and nonvascular plants, including mosses and green algae) (Nardmann *et al*., 2009; Van Der Graaff *et al*., 2009).

Some members of the WOX gene family have been shown to be involved in the regulation of AM. In *A. thaliana*, *WUS* is essential for the initiation and maintenance of AM (Wang *et al*., 2014a,b). Unlike the situation in *A. thaliana*, the AM of *O. sativa* is coregulated by *OsWUS* and *OsWOX4*. *OsWUS* is expressed only before meristem formation and not in the established AM, and *OsWOX4* is expressed only in the established AM, indicating that *OsWOX4* functions only in maintaining meristem activity (Ohmori *et al*., 2013; Lu *et al*., 2015; Tanaka *et al*., 2015). WOX9 and WOX11 are members of the intermediate clade and regulate the shoot meristem or AM. In the *A. thaliana wox9* mutant, the development of the embryo, apical meristem and root meristem is abnormal, and the growth and development of the axillary buds and roots is significantly inhibited (Skylar *et al*., 2010; Skylar & Wu, 2010). In addition, the loss of *WUS* expression in the *wox9* mutant indicates that WOX9 can positively regulate the expression of *WUS* (Wu *et al*., 2005). In *O. sativa*, *OsWOX9* (*Dwarftiller1*, *DWT1*) plays an important role in the development of rice tillers, and the *dwt1* mutant shows shorter tillers and a reduced tiller number (Wang *et al*., 2014c). In *A. thaliana* and *O. sativa*, *WOX11* mainly regulates the lateral root or crown root primordium (Liu *et al*., 2014; Hu & Xu, 2016). *wox11* mutants show crown root number and growth rate deficiencies, a dwarf phenotype and delayed flowering (Zhao *et al*., 2009). In crown and root development, OsWOX11 mediates the cytokinin pathway by inhibiting the expression of type-A *OsRR2*, thus enhancing cytokinin signalling to promote crown and root formation (Nardmann & Werr, 2006; Zhao *et al*., 2009). A recent study revealed that in addition to its function in crown root development, *OsWOX11* is also required for rice shoot development and can activate gene expression during the development of the rice shoot apical meristem by recruiting the H3K27me3 demethylase JMJ705 (Cheng *et al*., 2018).

On the basis of transcriptome data (accession number: SRP103184), we screened the expression of all annotated *WOX* genes during bulbil formation and identified two *WOX* genes closely related to bulbil formation, *LlWOX9* and *LlWOX11* (Fig. S1). In this study, our results showed that *LlWOX9* and *LlWOX11* were members of the intermediate clade and that their expression increased continuously during bulbil formation. The overexpression of *LlWOX9* and *LlWOX11* promoted bulbil formation, while the silencing of *LlWOX9* and *LlWOX11* inhibited bulbil formation, indicating that *LlWOX9* and *LlWOX11* are positive regulators of bulbil formation. Cytokinin type-B LlRRs can bind to the promoters of *LlWOX9* and *LlWOX11* to promote their transcription. In addition, LlWOX11 can enhance cytokinin signalling by inhibiting the transcription of type-A *LlRR9*. Our study enriches the understanding of the roles of the *WOX* gene family in regulating plant development. We also show for the first time that *WOX* genes cooperate with cytokinins to regulate the formation of bulbils. Our study lays a foundation for further research on the molecular mechanism of bulbil formation in lily.

## Materials and methods

### Plant materials and treatments

Bulbs of *Lilium lancifolium* of uniform size were harvested and buried in soil at 4°C at the Institute of Vegetables and Flowers, Chinese Academy of Agricultural Sciences (CAAS), Beijing, China, in November 2019. Well-grown stems with a height of 10 cm were selected according to an *in vitro* bulbil induction system (He *et al*., 2020), and stem segments were cultured on Murashige and Skong medium for bulbil induction. The stages of bulbil formation were divided into the bulbil initiation stage (S0–S2), bulbil primordium formation stage (S3–S4), and bulbil structure formation stage (S5) (He *et al*., 2020). Different stages of developing bulbils and different tissues (leaf axils at stage S4, shoot apex, leaf, stem, root, scale, stigma, ovary, anther and petal tissues) were collected for RNA extraction.

To determine whether *LlWOX9* and *LlWOX11* are immediately induced by cytokinins, 4 mM 6-BA was added to MS medium during bulbil formation, and stem segments at the S4 stage were treated with 10 mM 6-BA or with 0.05 mM NaOH as a control. Leaf axils were harvested at the S0-S5 stages and after 0, 0.5, 1.0, 1.5, 2.0 or 2.5 h of treatment.

### *Isolation of* LlWOX9 *and* LlWOX11 *genes and promoters*

According to our transcriptome data (accession number: SRP103184), we designed primers by using Primer 6 to clone the full-length sequences and promoters of *LlWOX9 and LlWOX11*. The full-length sequences of *LlWOX9 and LlWOX11* were cloned via RLM-RACE using the GeneRacer^TM^ Kit (Invitrogen, US) according to the kit protocol. To obtain the promoter sequences of *LlWOX9 and LlWOX11*, three gene-specific reverse primers were designed and a nested PCR program was used according to the protocol of a genome walking kit (Takara, Japan). The sequences of the primers used for amplification are shown in Table S1. Conserved protein domains were analyzed using SMART (http://smart.embl.de/). Phylogenetic analysis was performed using MEGA6 (http://mega6.software.informer.com/). Multiple sequence alignments were analysed using the DNAMAN software package. New PLACE (https://www.dna.affrc.go.jp/PLACE/?action=newplace) and PlantCARE (http://bioinformatics.psb.ugent.be/webtools/plantcare/html/) were used to analyse the *LlWOX9* and *LlWOX11* promoters.

### Real-time RT-PCR (qRT-PCR)

Total RNA from the different tissue and leaf axil specimens was extracted with an RNAprep Pure Plant Kit (TIANGEN, China) according to the kit protocol, and DNA contamination was removed with RNase-free DNase I. First-strand cDNA was synthesized with a Hifair^®^Ⅱ 1st Strand cDNA Synthesis Kit (gDNA digester plus) (Yeasen, China) according to the kit protocol. Gene-specific primers for qRT-PCR were designed with Primer 6.0 (Table S2). The *LilyActin* primer was used as an internal control (Xu *et al*., 2017), and SYBR^®^ Green Master Mix (No Rox) (Yeasen, Shanghai, China) was used in the reaction mixture according to the manufacturer’s instructions. qRT-PCR was conducted using the CFX96 Real-Time System (Bio-Rad, USA), with an initial denaturation step at 95°C for 3 min, followed by 40 cycles of denaturation at 95°C for 10 s, annealing at 60°C for 20 s, and extension at 72°C for 1 min. The 2^−ΔΔCt^ method was used to calculate the relative expression levels of the different genes (Livak & Schmittgen, 2001). Three biological and three technical replicates were performed to reduce error.

### Subcellular localization

The full-length cDNAs of *LlWOX9* and *LlWOX11* under the control of the 35S cauliflower mosaic virus promoter were cloned into the pCAMBIA 2300 vector using the pEASY^®^-Basic Seamless Cloning and Assembly Kit (Transgen Biotech, China). The sequences of the primer pairs used for amplification are shown in Table S3. The resulting plasmids were transferred into *Agrobacterium tumefaciens* strain GV3101. *Agrobacterium* cells were collected and suspended in infiltration buffer (10 mM methylester sulfonate, 10 mM MgCl_2_, and 150 mM acetosyringone, pH 5.7) at OD_600_ =0.8 and infiltrated into *Nicotiana benthamiana* leaves. 3 days after infiltration, the leaves were harvested and treated with 0.5 mg/ml DAPI (4ʹ,6-diamidino-2-phenylindole; Sigma). A Zeiss LSM 510 confocal scanning microscope was used to collect images.

### RNA fluorescence in situ hybridization

Leaf axils in S1-S5 were fixed with FAA, and after dehydration, clearing and embedding, paraffin sections of the leaf axils were sliced at a thickness of 0.8 μm. The obtained slides were rehydrated with xylene, digested with protease K (20 μg/mL) at 37°C, blocked with a 3% methanol-H2O2 solution for 25 min, with avidin (0.07%) at 37°C for 25 min and with biotin (0.005%) at 37°C for 15 min. Hybridization with the probes was performed at 37°C overnight in a moist chamber. After hybridization, the slides were washed in 2× SSC for 10 min, three times in 1× SSC for 5 min and once in 0.5× SSC for 10 min at 37°C. The avidin-labelled probe was detected with a streptavidin Alexa Fluor 405 conjugate (1:250) (Invitrogen). Antibodies were diluted in PBS containing 3% (w/v) BSA, and the slides were incubated with the antibodies for 30 min at 37°C. After antibody incubation, the slides were washed three times with 4× SSC containing 0.1% Tween 20, stained with 100 ng/mL DAPI in PBS for 30 min and dehydrated with ethanol. Confocal images were obtained using a Zeiss LSM 510 confocal scanning microscope.

### *Transformation of* A. thaliana *and* L. lancifolium

The full-length cDNAs of *LlWOX9* and *LlWOX11* were amplified by PCR and inserted into the pCAMBIA 3301 vector using the pEASY^®^-Basic Seamless Cloning and Assembly Kit (Transgen Biotech, China). All primers used are listed in Table S1. *A. thaliana* was transformed using *A. tumefaciens* strain GV3101 and the floral dip method (Clough & Bent, 1998).

Transgenic *A. thaliana* plants were selected on 1/2 Murashige and Skoog (MS) medium with 30 mg/L kanamycin. Transgenic *A. thaliana* plants were grown in climate-controlled boxes at 24°C under a 12/12 h light/dark cycle.

*L. lancifolium* was transformed using *A. tumefaciens* strain EHA105 via *Agrobacterium*-mediated vacuum infiltration. *Agrobacterium* cells were collected and suspended in infiltration buffer that contained 10 mM MgCl_2_, 200 mM acetosyringone and 10 mM MES (pH 5.6). The small stem segments of *L. lancifolium* were submerged in infiltration solution and then subjected to -50 kPa vacuum for 10 min. The infiltrated segments were washed with distilled water three times and then were grown on MS medium with 30 g/L sucrose and 6 g/L agar (pH 5.8) in the dark at 20°C for 1 d, followed by growth at 22°C under a 16/8 h light/dark cycle. The rate of bulbil formation was assessed after one week of culture, and RNA was extracted from leaf axils to measure the expression of the target genes. Each treatment consisted of three experimental replicates, with 30 leaf axils per replicate.

### Virus-induced gene silencing (VIGS)

For the generation of pTRV2-LlWOX9 and pTRV2-LlWOX11, gene-specific fragments of ∼300 bp were cloned into the pTRV2 vector using the pEASY^®^-Basic Seamless Cloning and Assembly Kit (Transgen Biotech, China). Five pTRV2-LlRR vectors were constructed as previously described (He *et al*., 2021). The primer pairs used to generate the TRV vectors are shown in Table S3. VIGS was performed using *A. tumefaciens* strain EHA105 and vacuum infiltration method (He *et al*., 2021). The rate of bulbil formation was assessed after two weeks of culture, and RNA was extracted from leaf axils to measure the expression of the target genes. Each treatment consisted of three experimental replicates, with 30 leaf axils per replicate.

### GUS staining

A 1318 bp fragment upstream of the start codon of *LlWOX9* and a 2351 bp fragment upstream of the start codon of *LlWOX11* were introduced into the pCAMBIA 3301 vector, and the *35S* promoter was replaced using the pEASY®-Basic Seamless Cloning and Assembly Kit (Transgen Biotech, China). The constructed plasmids were transferred into *A. tumefaciens* strain EHA105. The method of *N. benthamiana* leaf infiltration was the same as that used in the subcellular localization assay. Stem segments at S0 and S5 were used for vacuum infiltration according to the method described above. Three days after infiltration, the leaves and stem segments were harvested and treated with GUS staining solution (Solarbio, China) according to the kit protocol. After staining, the leaves and stem segments were washed and cleared with 70% ethanol for more than 24 h before image capture using a Leica Microsystems DM5500B instrument (Wetzlar, Germany).

### Yeast one-hybrid assay

Y1H analysis was performed according to the method described by Lin *et al*. (2013). Briefly, the full-length coding regions of five *LlRR*s and *LlWOX11* were cloned into the pGADT7 vector to generate the pGADT7-LlRRs and pGADT7-LlWOX11 constructs. Various truncated versions of the promoter regions of *LlWOX9* and *LlWOX11* were amplified and ligated into the pABAi reporter vector. The constructs were then cotransformed into the yeast strain EGY48. Transformants were grown on SD-Trp/-Ura plates for 3 d at 28°C. The interactions were determined based on the growth ability of the cotransformants on medium supplemented with aureobasidin A (AbA).

### Dual-luciferase reporter assay

The coding sequence of *LlWOX11* was cloned into the pCAMBIA 3301 vector using the pEASY^®^-Basic Seamless Cloning and Assembly Kit (Transgen Biotech, China). Five pCAMBIA 3301-LlRR vectors were constructed as previously described (He *et al*., 2021). A 1318 bp fragment upstream of the start codon of *LlWOX9* and a 2351 bp fragment upstream of the start codon of *LlWOX11* were introduced into the pluc-35Rluc vector using the pEASY^®^-Basic Seamless Cloning and Assembly Kit (Transgen Biotech, China). The primers used to generate the constructs are listed in Table S3. The constructed plasmids were transformed into *A. tumefaciens* strain GV3101. Different effectors were subsequently coinfiltrated with the reporter into *N. benthamiana* leaves using a syringe. At 3 d after infiltration, 2-cm-diameter leaf discs were harvested and ground in liquid nitrogen. The activities of firefly and Renilla luciferase were measured with a Dual-Luciferase Reporter Assay System (Promega) using a GloMax 20/20 luminometer (Promega).

### EMSAs

To construct plasmids for the expression of the recombinant LlWOX11 protein in *Escherichia coli*, the full-length cDNA was amplified and cloned into the pMal-c2X vector, which was expressed in the *Escherichia coli* strain BL21 (DE3) cell line. The pET32a-LlRR1 vector was constructed as previously described (He *et al*., 2021). The primers are listed in Table S3. Protein expression was induced by incubation in 1 mM IPTG at 16°C at 160 rpm for 24 h. Protein purification was carried out using an amylose resin purification system (NEB) following the manufacturer’s instructions. Double-stranded oligonucleotide probes were synthesized and labelled with biotin at the 5′end. EMSA was carried out using the LightShift^®^ Chemiluminescent EMSA Kit (Thermo Fisher Scientific, USA). Competition experiments were performed with different amounts of nonlabelled oligonucleotides. The mutated competitors were generated by replacing eight base pairs in the WOX binding elements (TTAATGAG to AAAAAAAA).

## Results

### *Full-length cloning and sequence analysis of* LlWOX9 *and* LlWOX11

On the basis of transcriptome data (accession number: SRP103184), we cloned the full-length sequences of *LlWOX9* (1008 bp) and *LlWOX11* (699 bp) by RLM-RACE and found that they encoded 335 and 232 amino acids, respectively (**Fig. 1a**). Amino acid sequence analysis showed that both LlWOX9 and LlWOX11 contained HOX domains at the N-terminus (**Fig. 1a**). Sequence alignment confirmed a conserved HOX domain at the N-terminus in LlWOX9 and LlWOX11 (**Fig. 1b,c**). A phylogenetic tree of LlWOX9, LlWOX11 and the members of the WOX transcription factor family in *A. thaliana* was constructed, and the results showed that LlWOX9 and LlWOX11 belonged to the intermediate evolutionary branch of the WOX family (**Fig. 1d**). Phylogenetic tree of WOX9 and WOX11 from different species showed that LlWOX9 and LlWOX11 were clustered with the sequences of other monocotyledonous species and were closely related to the WOX9 and WOX11 amino acid sequences of *Palmaceae* plants (**Fig. 1e, f**).

**Fig. 1.**
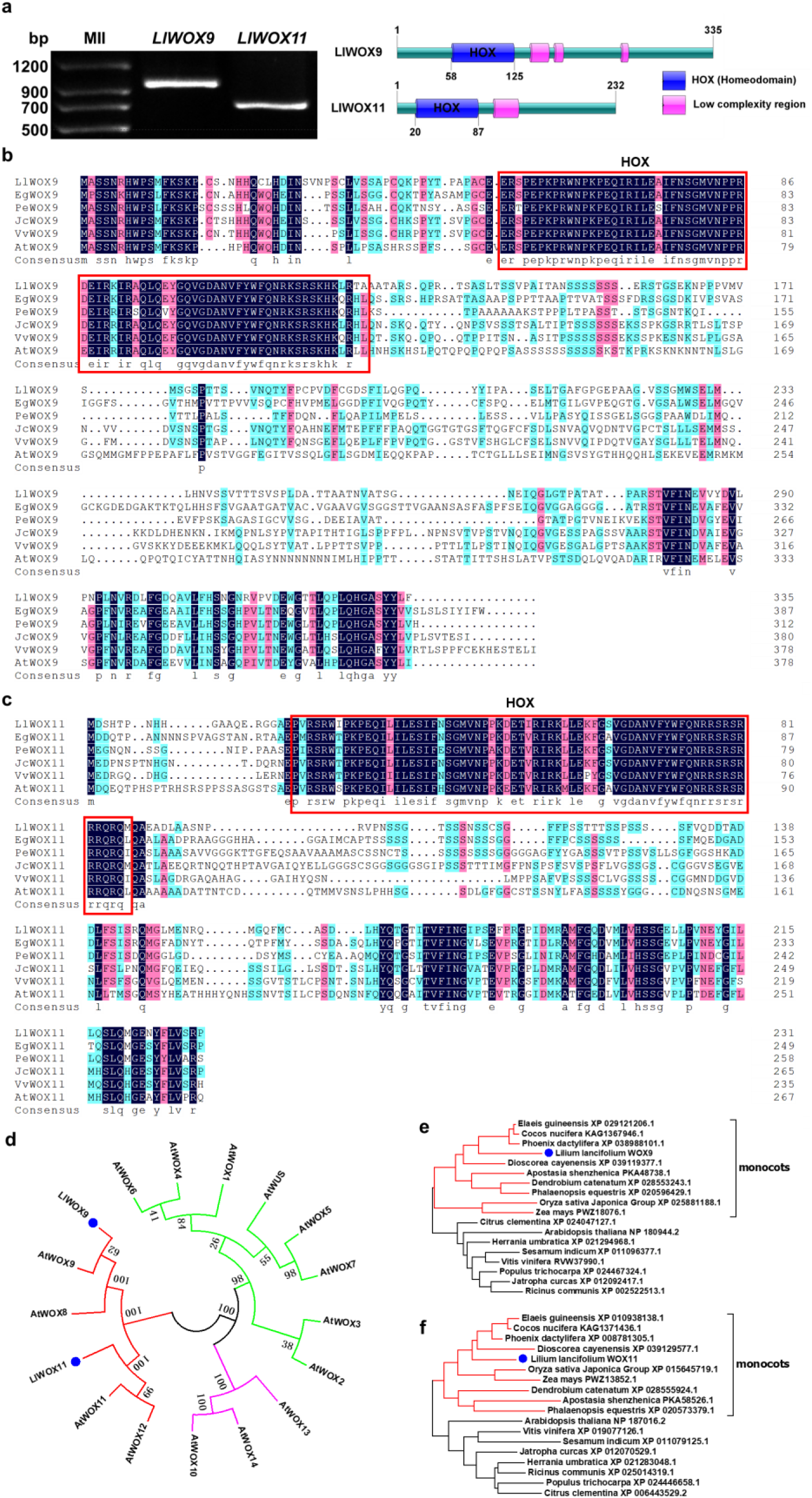
Full-length cloning, sequence alignment and phylogenetic tree of *LlWOX9* and *LlWOX11*. **a**: Full-length cloning and domain prediction of *LlWOX9* and *LlWOX11*. **b**: Multiple sequence alignment of LlWOX9 with sequences of other species. **c**: Multiple sequence alignment of LlWOX11 with sequences of other species. The red boxes in B and C represent the HOX domain. *Ll*: *Lilium lancifolium*, *Eg*: *Elaeis guineensis* (EgWOX9, XP 029121206.1; EgWOX11, XP 010938138.1), *Pe*: *Phalaenopsis equestris* (PeWOX9, XP 020596429.1; PeWOX11, XP 020573379.1), *Jc*: *Jatropha curcas* (JcWOX9, XP 012092417.1; JcWOX11, XP 012070529.1), *Vv*: *Vitis vinifera* (VvWOX9, RVW37990.1; VvWOX11, XP 019077126.1), *At*: *Arabidopsis thaliana* (AtWOX9, NP 180994.2; AtWOX11, NP 187016.2). **d**: Neighbour-joining tree of the LlWOX9 and LlWOX11 amino acid sequences of *L. lancifolium* and WOX family amino acid sequences from *A. thaliana*. **e**: Neighbour-joining tree of the LlWOX9 amino acid sequence of *L. lancifolium* and WOX9 amino acid sequences from other species. **f**: Neighbour-joining tree of the LlWOX11 amino acid sequence of *L. lancifolium* and WOX11 amino acid sequences from other species. Bootstrap values from 1,000 replicates were used to assess the robustness of the tree.

### *Expression pattern and subcellular localization of* LlWOX9 *and* LlWOX11

To study the subcellular localization of LlWOX9 and LlWOX11, we fused the LlWOX9 and LlWOX11 proteins with a green fluorescent protein (GFP) tag and introduced them into the leaves of *Nicotiana benthamiana*. The subcellular localization results showed that the GFP signals of the LlWOX9-GFP and LlWOX11-GFP fusion proteins were located in the nuclei of tobacco leaf epidermal cells (**Fig. 2a**), indicating that LlWOX9 and LlWOX11 may function as transcription factors in the nucleus.

**Fig. 2.**
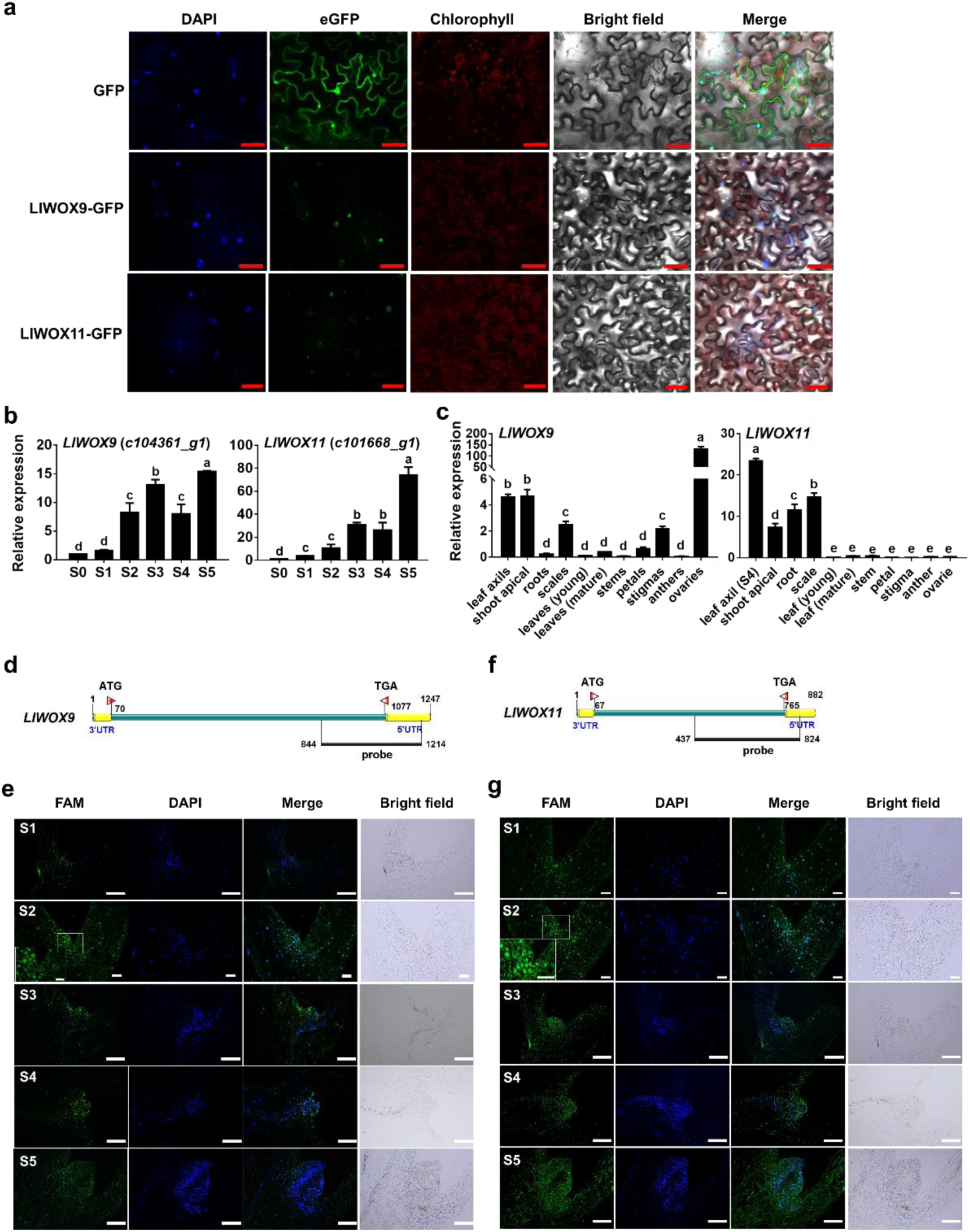
Subcellular localization, expression patterns and fluorescence in situ hybridization of *LlWOX9* and *LlWOX11*. **a**: Subcellular localization of LlWOX9-GFP and LlWOX11-GFP proteins in *Nicotiana benthamiana* leaf epidermal cells with 4ʹ,6-diamidino-2-phenylindole (DAPI) staining. Scale bars = 50 μm. **b**: *LlWOX9* and *LlWOX11* expression during bulbil formation. **c**: *LlWOX9* and *LlWOX11* expression in different tissues. Values are means ± SDs (n=3). Lowercase letters (a-d in B; a-e in C) indicate statistically significant differences at *P* < 0.05. **d**: Gene-specific probe of *LlWOX9* used in fluorescence in situ hybridization. **e**: Fluorescence in situ hybridization of *LlWOX9* during bulbil formation. **f**: Gene-specific probe of *LlWOX11* used in fluorescence in situ hybridization. **g**: Fluorescence in situ hybridization of *LlWOX11* during bulbil formation. Scale bar in A (S2) and B (S1, S2), 100 μm. Scale bar in A (S1, S3-S5) and B (S3-S5), 500 μm.

The expression of *LlWOX9* and *LlWOX11* increased continuously during bulbil formation (**Fig. 2b**). *LlWOX9* was mainly expressed in the leaf axil (S4 stage), shoot apical meristem, scale, stigma and ovary, with the highest relative expression in the ovary and the second highest in the leaf axil (**Fig. 2c**). *LlWOX11* was mainly expressed in the leaf axil (S4 stage), shoot apical tissue, root and scale, and the highest relative expression was found in the leaf axil (S4 stage) (**Fig. 2c**). The relatively high expression of *LlWOX9* and *LlWOX11* in leaf axils further indicated that *LlWOX9* and *LlWOX11* might be involved in bulbil formation.

We further detected the expression of *LlWOX9* and *LlWOX11* during bulbil formation by fluorescence in situ hybridization (FISH). Gene-specific sequences containing the 3’-UTRs of *LlWOX9* and *LlWOX11* were selected to synthesize the FAM -labelled fluorescent probes (**Fig. 2d,f**). Our results showed that although the *LlWOX9* and *LlWOX11* fluorescent signals could be detected throughout the analysed tissue, the fluorescent signals of *LlWOX9* and *LlWOX11* were mainly located in the leaf axil and gradually increased during bulbil formation (**Fig. 2e,g**). In addition, the fluorescent signals of *LlWOX9* and *LlWOX11* appeared on a differentiated scale (S5 stage) (**Fig. 2e,g**). These results further indicated that *LlWOX9* and *LlWOX11* are involved not only in the formation of the bulbil primordium but also in the differentiation of the bulbil scale.

### *Overexpression of* LlWOX9 *and* LlWOX11 *increases the number of branches in* A. thaliana

Bulbils can be considered a special type of branch. To investigate the function of *LlWOX9* and *LlWOX11* in *A. thaliana* branches, we generated transgenic *A. thaliana* lines. The transgenic lines were identified using the 35S-F and LlWOX9-R or LlWOX11-R primers. An ∼1200 or ∼800 bp band was amplified from the genomic DNA of the transgenic lines, and no corresponding bands were amplified from control plants (**Fig. 3a**). Our results demonstrated that overexpression of *LlWOX9* or *LlWOX11* in *A. thaliana* increased the number of branches and promoted the formation of accessory buds on inflorescences (**Fig. 3b,d**). The number of branches was significantly higher in the *35S::LlWOX9* and *35S::LlWOX11* transgenic lines than in the wild type (**Fig. 3c,e**). Interestingly, we found that the *35S::LlWOX9* transgenic lines showed some abnormal phenotypes, such as the development of the inflorescence branches into a single flower and the abnormal elongation of stem internodes in rosette leaves (**Fig. 3b**).

**Fig. 3.**
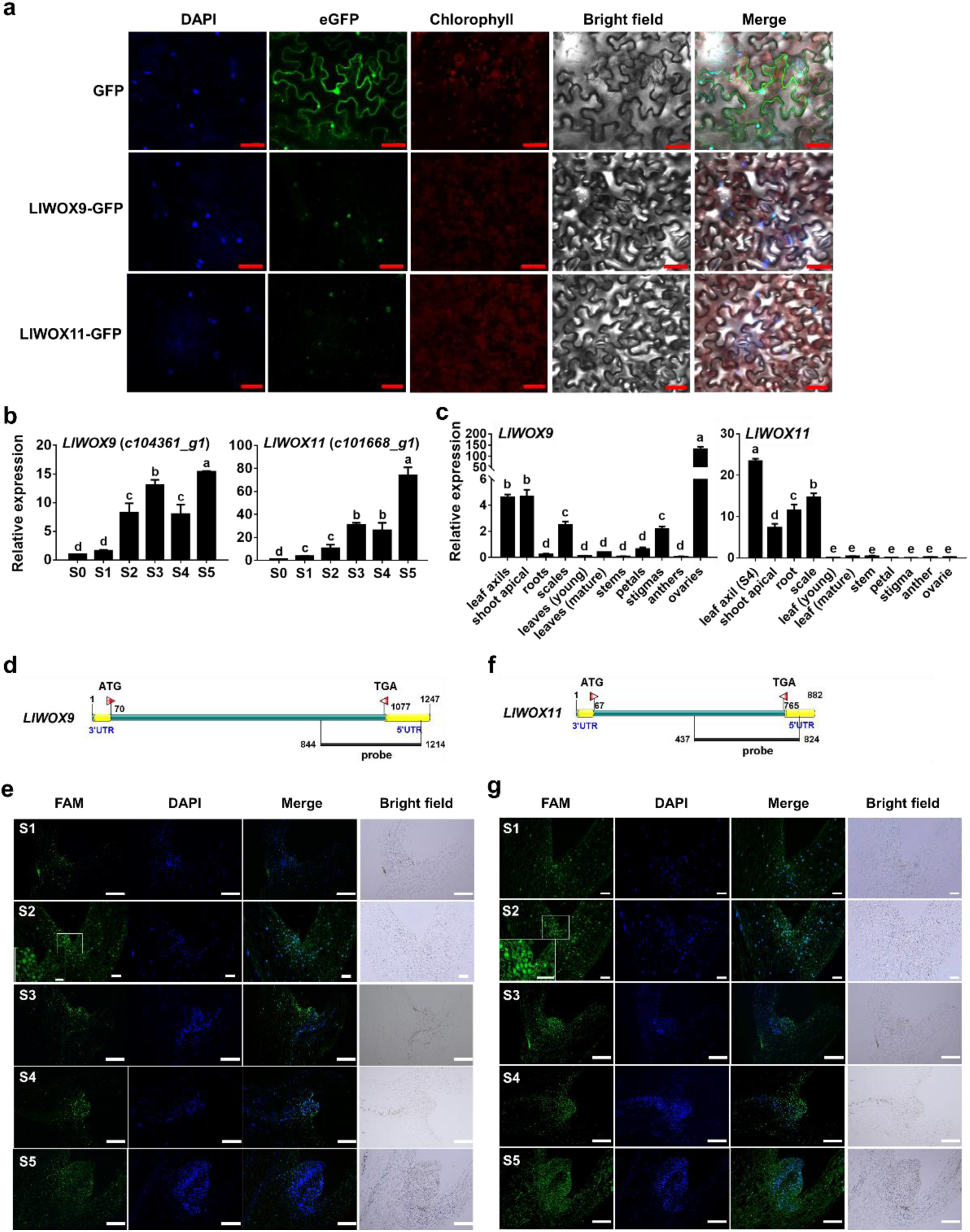
The phenotypes of *35S::LlWOX9* and *35S::LlWOX11* transgenic lines and wild-type *Arabidopsis thaliana* plants. **a**: The transgenic plants of the T3 generation of *A. thaliana* were detected by PCR. ‘+’ indicates the positive control and ‘-’ indicates the negative control. 1-5 represent different transgenic lines overexpressing *LlWOX9*, 6-11 represent different transgenic lines overexpressing *LlWOX11*. **b**: The branching phenotypes of wild-type Col and transgenic plants overexpressing *LlWOX9*. **c**: The numbers of branches on wild-type Col and transgenic plants overexpressing *LlWOX9*. **d**: The branching phenotypes of wild-type Col and transgenic plants overexpressing *LlWOX11*. **e**: The numbers of branches on wild-type Col and transgenic plants overexpressing *LlWOX11*.

### LlWOX9 *and* LlWOX11 *overexpression promotes bulbil formation*

To preliminarily understand the functions of *LlWOX9* and *LlWOX11* during bulbil formation, we further evaluated the functions of *LlWOX9* and *LlWOX11* via their transient overexpression in leaf axils through an *in vitro* bulbil induction system. Our results showed that after 6 d of culture, most of the developing leaf axils in the control group and the *35S::GUS* treatment group were still in the S3 stage (**Fig. 4a**), but the overexpression of *LlWOX9* and *LlWOX11* could significantly promote the formation of bulbils (**Fig. 4a**), and the rate of bulbil induction was significantly higher than that in the control group and the *35S::GUS* treatment group (**Fig. 4b**). These results indicated that *LlWOX9* and *LlWOX11* play important roles during bulbil formation.

**Fig. 4.**
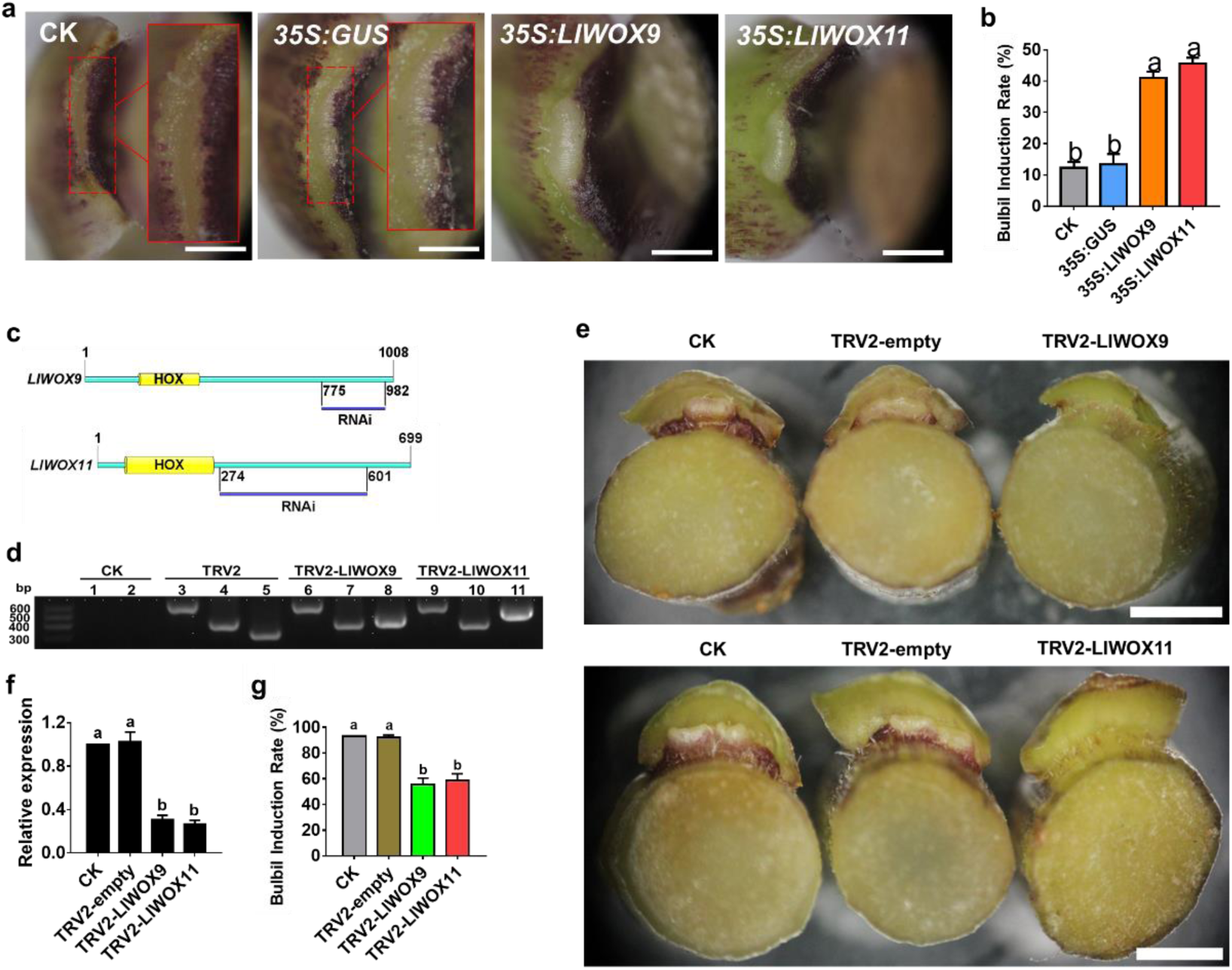
Phenotype and relative expression of *LlWOX9* and *LlWOX11* in leaf axils after overexpressing or silencing *LlWOX9* and *LlWOX11*. **a**: The phenotype of the leaf axil after the transient overexpression of *LlWOX9* and *LlWOX11*. **b**: The bulbil induction rate after the transient overexpression of *LlWOX9* and *LlWOX11*. The red box in figure A shows an enlargement of the indicated portion of the leaf axil. Values are means ± SDs (n=3). Scale bar in A, 1 mm. **c**: Specific fragments of genes used in VIGS experiments. **d**: PCR was used to detect the presence of the TRV1 and TRV2 viruses in the leaf axils. CK is the negative control, TRV2 is the positive control. Lanes 1, 3, 6 and 9 show TRV1 detection; 2, 4, 7 and 10 show the detection of coat proteins in TRV2; and lanes 5, 8 and 11 show the detection of inserts in TRV2. **e**: The phenotype of the leaf axil after silencing *LlWOX9* and *LlWOX11*. **f**: The relative expression of *LlWOX9* and *LlWOX11* in leaf axils after silencing *LlWOX9* and *LlWOX11*. **g**: The bulbil induction rate after silencing *LlWOX9* and *LlWOX11*. Values are means ± SDs (n=3). Scale bar in C, 50 mm. Lowercase letters (a-b in D, E) indicate statistically significant differences at *P* < 0.05.

### LlWOX9 *and* LlWOX11 *silencing inhibits bulbil formation*

To further understand the functions of *LlWOX9* and *LlWOX11* during bulbil formation, we constructed the TRV2-LlWOX9 and TRV2-LlWOX11 silencing vectors by selecting specific fragments of the *LlWOX9* and *LlWOX11* genes (**Fig. 4c**). After 12 d of infection with the empty TRV2 vector and the recombinant TRV2-LlWOX9 or TRV2-LlWOX11 vector, leaf axil cDNAs were obtained, and TRV1-F/R and TRV2-F/R were used for PCR-based detection. The results showed that in leaf axils infected with the empty TRV2 vector, TRV2-LlWOX9 or TRV2-LlWOX11, the target bands of pTRV1, the coat protein in pTRV2 and the insert fragment in pTRV2 could be detected (**Fig. 4d**). These results indicated that TRV2, TRV2-LlWOX9 and TRV2-LlWOX11 were successfully inserted and expressed in the genome of *L. lancifolium*.

The silencing of the *LlWOX9* and *LlWOX11* genes in leaf axils was detected by qRT-PCR. The results showed that the expression of *LlWOX9* and *LlWOX11* in leaf axils infected with TRV2-LlWOX9 or TRV2-LlWOX11 was significantly lower than that in the control and the leaf axils infected with TRV2 (**Fig. 4f**). These findings indicated that *LlWOX9* and *LlWOX11* were effectively silenced in TRV2-LlWOX9- and TRV2-LlWOX11-infected leaf axils, respectively.

The silencing experiment results showed that after *LlWOX9* and *LlWOX11* silencing, the formation of bulbils was inhibited compared to that in the control group and the empty TRV2 treatment group (**Fig. 4e**) and the rate of bulbil induction decreased significantly (**Fig. 4g**). These results indicated that *LlWOX9* and *LlWOX11* play important roles by positively regulating bulbil formation.

### *Cytokinins induce the expression of* LlWOX9 *and* LlWOX11

Our previous studies have revealed that cytokinins can promote bulbil formation through type-B RRs. To study whether the expression of *LlWOX9* and *LlWOX11* is regulated by cytokinins, we detected the expression of *LlWOX9* and *LlWOX11* after exogenous cytokinin treatment and the silencing of type-B *LlRR*s. The results showed that after treatment with 6-BA, the expression of *LlWOX9* and *LlWOX11* during bulbil formation was significantly higher than in the control group (**Fig. 5a**). To further study whether the expression of *LlWOX9* and *LlWOX11* was directly induced by exogenous cytokinins, we treated leaf axils at the S4 stage with 6-BA. The results showed that after exogenous 6-BA treatment, the expression of *LlWOX9* and *LlWOX11* was rapidly induced (**Fig. 5b**), indicating that exogenous cytokinins could induce the expression of *LlWOX9* and *LlWOX11*. In addition, because cytokinins regulate downstream genes through type-B RRs, we detected the expression of *LlWOX9* and *LlWOX11* after the silencing of five type-B *LlRR*s in leaf axils. The results showed that the expression of *LlWOX9* and *LlWOX11* decreased significantly after the silencing of a single type-B *LlRR* gene, while after the silencing of five type-B *LlRR*s, the relative expression of *LlWOX9* and *LlWOX11* was almost undetectable (**Fig. 5c**). These results suggest that cytokinins can induce the expression of *LlWOX9* and *LlWOX11* and that type-B LlRRs may directly regulate the expression of *LlWOX9* and *LlWOX11*.

**Fig. 5.**
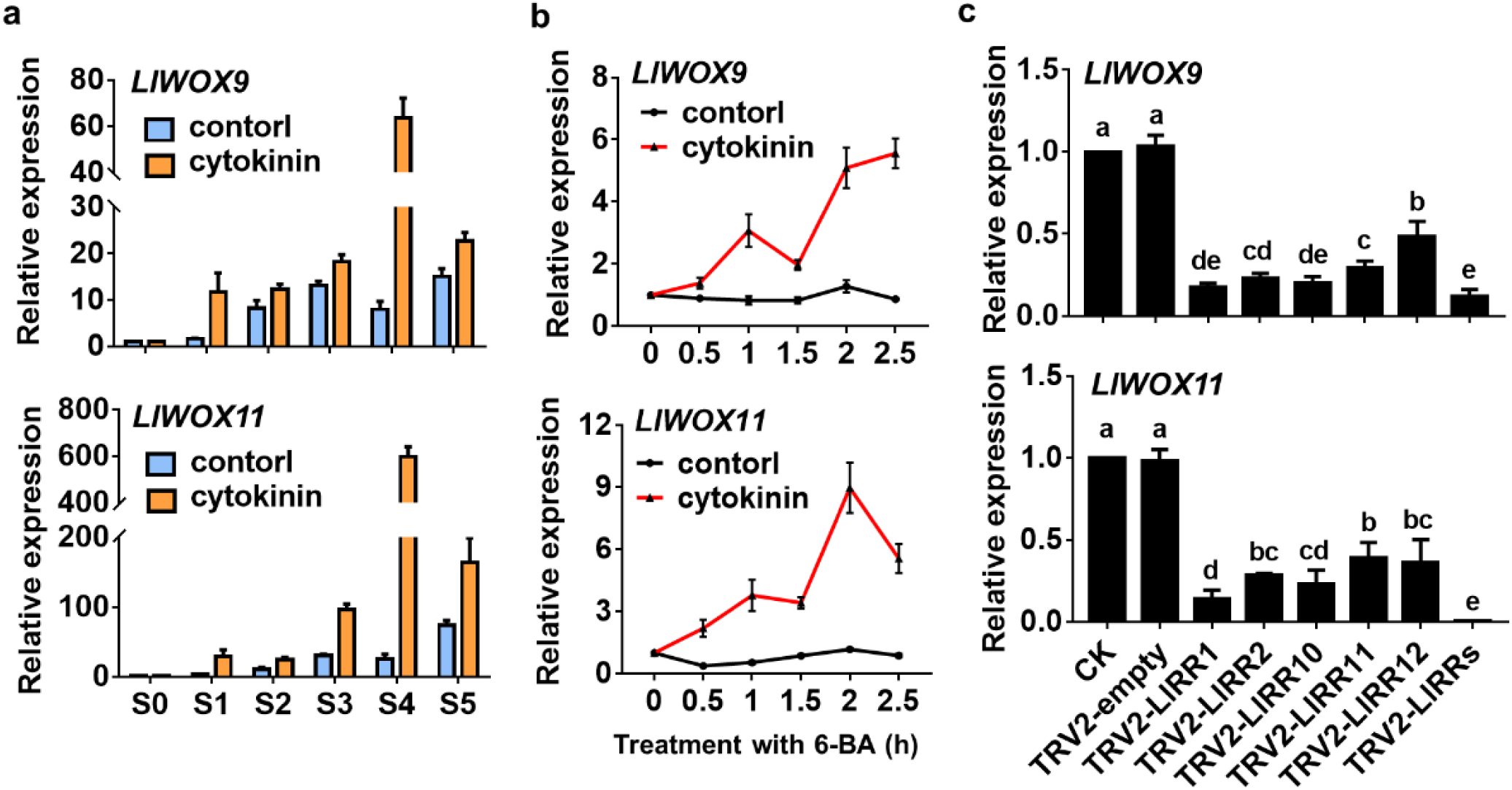
Expression of *LlWOX9* and *LlWOX11* after treatment with 6-BA and type-B *LlRR* silencing. **a**: Expression of *LlWOX9* and *LlWOX11* after treatment with 4 μM 6-BA during bulbil formation. **b**: Expression of *LlWOX9* and *LlWOX11* in leaf axils at stage S4 after 10 mM 6-BA treatment. **c**: Expression of *LlWOX9* and *LlWOX11* after type-B *LlRR* silencing. Lowercase letters (a-e in C) indicate statistically significant differences at *P* < 0.05.

### *Type-B LlRRs promote the transcription of* LlWOX9 *and* LlWOX11

To study whether type-B LlRRs regulate the transcription of *LlWOX9* and *LlWOX11*, we first cloned the promoter sequences of *LlWOX9* and *LlWOX11* via the chromosome walking technique and obtained promoters with lengths of 1318 bp and 2351 bp, respectively (**Fig. 6a**). The use of the New Place and PlantCARE online element prediction tools showed that the promoters of *LlWOX9* and *LlWOX11* contained a large number of type-B RR binding elements (NGATT/C) (**Fig. 6a**). Furthermore, we studied the promoter activities of *LlWOX9* and *LlWOX11* in tobacco leaves and *L. lancifolium* stem segments. GUS staining results showed that both the *LlWOX9* and *LlWOX11* promoters were active and that their activity was weaker than that of the 35S promoter (**Fig. 6b**). In the stem segments, we observed stronger GUS staining of the promoters of *LlWOX9* and *LlWOX11* in the leaf axils (**Fig. 6b**), indicating that the promoters of *LlWOX9* and *LlWOX11* may show tissue specificity.

**Fig. 6.**
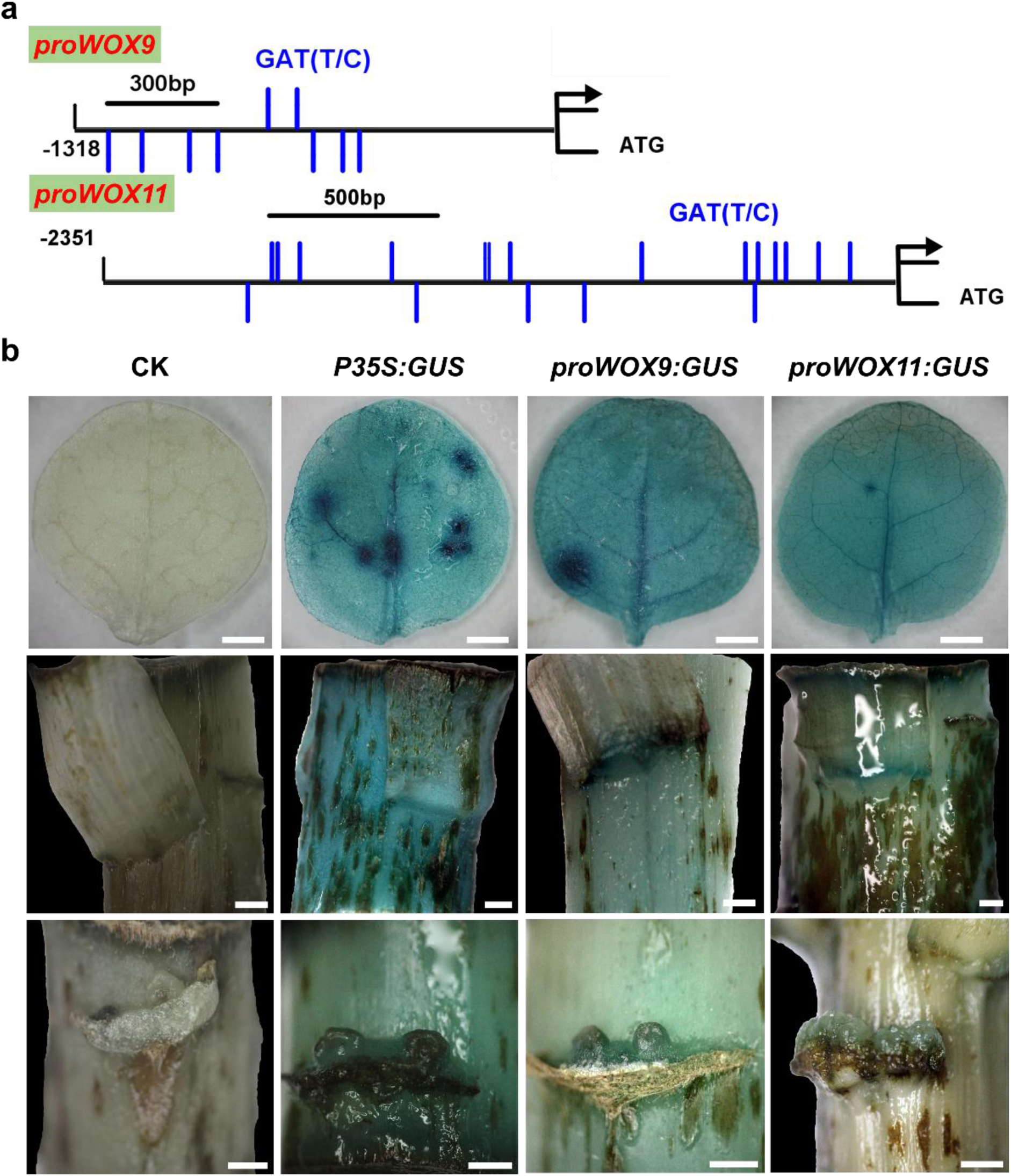
Detection of *LlWOX9* and *LlWOX11* promoter activity. **a**: The *LlWOX9* and *LlWOX11* promoters contain a large number of type-B RR binding elements. **b**: The expression of *GUS* was driven by the *35S*, *LlWOX9* and *LlWOX11* promoters, and GUS staining was performed in *Nicotiana benthamiana* leaves and *Lilium lancifolium* stems. In B, the *N. benthamiana* leaf scale bar is 50 mm, and the *L. lancifolium* stem scale bar is 1 mm.

Then, we divided the *LlWOX9* and *LlWOX11* promoters into three or five segments, respectively, according to the positions of GATT/C elements to construct yeast bait vectors (**Fig. 7a**). Yeast one-hybrid results showed that five type-B LlRRs could strongly bind the promoter sequences of *LlWOX9* and *LlWOX11*. Among these sequences, the *proWOX9-I* fragment could be bound by LlRR1, LlRR2, LlRR10 and LlRR12 (**Fig. 7b**); the *proWOX9-II* fragment could be bound by LlRR2, LlRR11 and LlRR12 (**Fig. 7b**); and the *proWOX9-III* fragment could not be bound by any LlRR because it contained no predicted binding element (**Fig. 7b**). The *proWOX11-I* fragments could be bound by LlRR1, LlRR10 and LlRR12 (**Fig. 7c**); the *proWOX11-II* fragments could be bound by LlRR1, LlRR2 and LlRR10 (**Fig. 7c**); the *proWOX11-III* fragments could be bound by LlRR1, LlRR10, LlRR11 and LlRR12 (**Fig. 7c**); the *proWOX11-IV* fragments could be bound by LlRR2, LlRR10 and LlRR11 (**Fig. 7c**); and the *proWOX11-V* fragments could be bound by LlRR1, LlRR2 and LlRR12 (**Fig. 7c**).

**Fig. 7.**
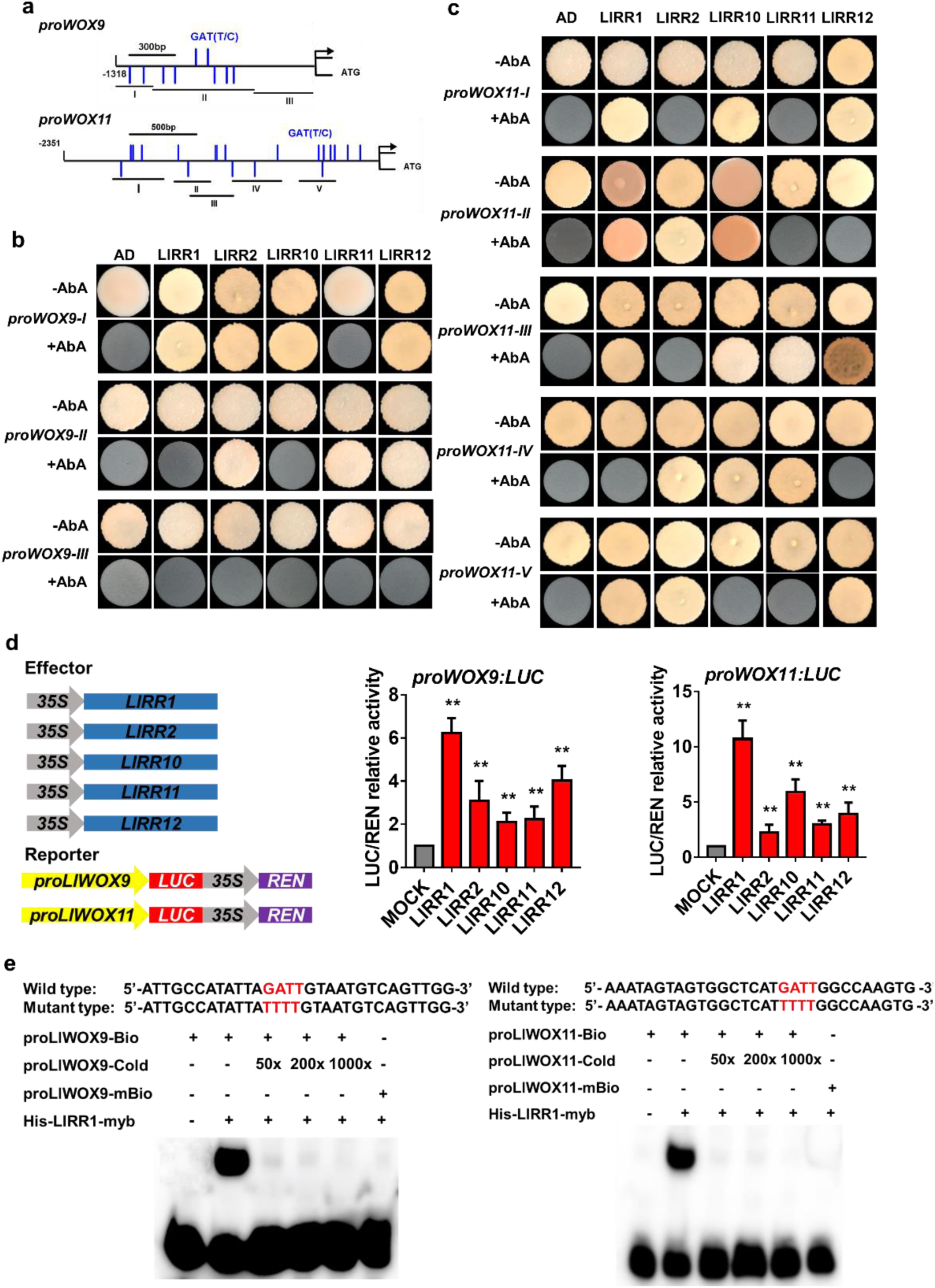
Yeast one-hybrid and dual-luciferase reporter assays and EMSAs of type-B LlRRs with the *LlWOX9* and *LlWOX11* promoters. **a**: Division of the *LlWOX9* and *LlWOX11* promoters into fragments according to the location of type-B RR binding elements (GATT/C). **b**: Yeast one-hybrid assays between five type-B LlRRs and *LlWOX9* promoter fragments. **c**: Yeast one-hybrid assays between type-B LlRRs and *LlWOX11* promoter fragments. **d**: The transient activation test in tobacco leaves verified the transcriptional activation ability of the five type-B LlRRs toward the *LlWOX9* and *LlWOX11* promoters. **e**: The binding ability of His-LlRR1 protein toward the *proLlWOX9-1* and *proLlWOX11-2* fragments was verified by EMSAs. The binding element GATT was mutated to TTTT in the mutant probe. Asterisks in A indicate significant differences compared with the control, with two asterisks indicating *P* < 0.01.

Furthermore, we studied the transcriptional activation ability of five type-B LlRRs toward the *LlWOX9* and *LlWOX11* promoters in tobacco leaves. The results showed that compared with the control group, all five type-B LlRRs could significantly activate the transcription of the *LlWOX9* and *LlWOX11* promoters (**Fig. 7d**). In addition, we selected 30 bp fragments of the *proWOX9-I* and *proWOX11-II* fragments containing GATT/C elements to synthesize biotin-labelled probes for electrophoretic mobility shift assays (EMSAs). The results showed that the His-LlRR1 protein could directly bind the *proWOX9-I* and *proWOX11-II* fragments (**Fig. 7e**).

### LlWOX11 *mediates the cytokinin pathway by inhibiting the transcription of* LlRR9

*LlRR9* is a type-A response regulator gene whose product is a negative regulator of cytokinin signalling. Our previous studies have revealed that LlRR9 is involved in bulbil formation and transcriptional regulation by LlRR1. In this study, we found a WOX binding element (TTAATGAG) 2097 bp upstream of ATG in the promoter of *LlRR9* (**Fig. 8a**). To determine whether *LlRR9* is a downstream gene directly regulated by LlWOX9 or LlWOX11, we carried out dual-luciferase reporter and yeast one-hybrid assays and EMSAs. Our results showed that LlWOX9 did not affect transcription from the *LlRR9* promoter (data not shown), but LlWOX11significantly inhibited transcription from the *LlRR9* promoter (**Fig. 8b**). The results of yeast one-hybrid assays showed that LlWOX11 could bind the *LlRR9* promoter fragment containing the TTAATGAG element (**Fig. 8c**), while LlWOX9 did not show any binding capacity (data not shown). Furthermore, a biotin-labelled probe was synthesized by selecting a 30 bp fragment of the *LlRR9* promoter containing the TTAATGAG element for EMSA. The EMSA results showed that LlWOX11 could directly bind the *LlRR9* promoter sequence (**Fig. 8d**).

**Fig. 8.**
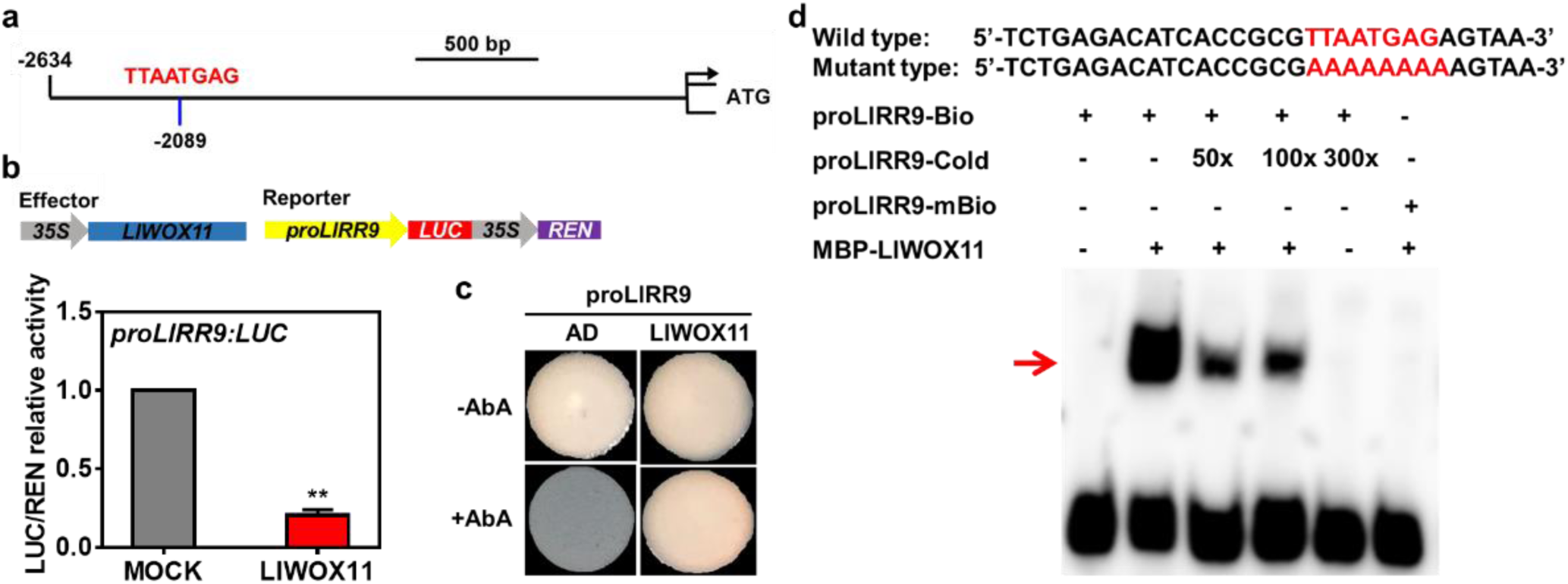
The interaction between LlWOX11 and the *LlRR9* promoter was verified by dual-luciferase reporter and yeast one-hybrid assays and EMSA. **a**: The transient activation test in tobacco leaves verified the transcriptional activation ability of LlWOX11 toward the *LlRR9* promoter. **b**: The binding ability of LlWOX11 toward the *LlRR9* promoter was verified by a yeast one-hybrid assay. **c**: The binding ability of the MBP-LlWOX11 protein toward the *proLlRR9* fragment was verified by EMSA. The binding element TTAATGA was mutated to AAAAAAA in the mutant probe. **d**: The transient activation test in tobacco leaves verified the transcriptional activation ability of LlWOX11 toward the *LlWOX9* promoter. Asterisks in C indicate significant differences compared with the control, with two asterisks indicating *P* < 0.01.

## Discussion

### LlWOX9 *and* LlWOX11 *are widely expressed in* L. lancifolium

*WOX9* and *WOX11* show broad expression profiles in different species (Wu *et al*., 2005; Zhao *et al*., 2009; Cheng *et al*., 2014; Li *et al*., 2018). In this study, we showed that *LlWOX9* was mainly expressed in the leaf axil (S4 stage), shoot apical tissue, scale, stigma and ovary and that *LlWOX11* was mainly expressed in the leaf axil (S4 stage), shoot apical tissue, root and scale. Through in situ hybridization and the analysis of GUS reporter promoter fusions, *WOX9* could be detected in the vegetative SAM, leaf primordia, floral meristems, early floral organs and root meristematic zone (Wu *et al*., 2005). *OsWOX11* mRNA was detected in calli, roots, 7-d-old seedlings, SAM, leaf primordia, and young leaves (Zhao *et al*., 2009). Our results similarly showed that *LlWOX9* and *LlWOX11* were highly expressed in leaf axils (S4 stage) and apical shoots. In accord with the reported relationships of *WOX11* with the development of lateral roots and crown roots in *A. thaliana* and *O. sativa* (Liu *et al*., 2014; Hu & Xu, 2016), our results showed that *LlWOX11* was highly expressed in roots. Unlike previous research results, our results showed that *LlWOX9* presented the highest relative expression in female reproductive organs but was almost undetectable in male reproductive organs. In *M. truncatula*, *MtWOX9* is mainly expressed in nodules, leaves and flowers but is expressed at lower levels in ovaries (Li *et al*., 2018). *OsWOX9* is expressed in axillary tillers, panicles, stamen primordia and pistil primordia but can only be detected in anthers after flower development maturity (Cheng *et al*., 2014).

### LlWOX9 *and* LlWOX11 *are positive regulators during bulbil formation*

Members of the intermediate clade are widely expressed in plants and usually play a role in maintaining meristem cell division (Wu *et al*., 2007; Breuninger *et al*., 2008; Zhao *et al*., 2009). In *A. thaliana*, when *WOX9* function is lost, cells divide abnormally, and the development of the shoot meristem is defective (Wu *et al*., 2005; Skylar *et al*., 2010). In the *Oswox11* mutant, the crown root number and plant height are decreased, and the growth rate and flowering time are delayed, while in *OsWOX11*-overexpressing lines, the number of crown roots is increased, ectopic crown roots form at the base of the spikelets, and the growth rate increases significantly (Zhao *et al*., 2009). In this study, the overexpression of *LlWOX9* and *LlWOX11* in *A. thaliana* significantly increased branch numbers and promoted bulbil formation in *L. lancifolium*. We found that the rate of bulbil induction decreased after the silencing of *LlWOX9* and *LlWOX11* expression but increased significantly after *LlWOX9* and *LlWOX11* overexpression, indicating that *LlWOX9* and *LlWOX11* are positive regulators of bulbil formation. After the overexpression of *LlWOX9* and *LlWOX11*, we also observed the abnormal proliferation of axillary tissue cells and the development of large purple-black bulbils on leaf axils (Fig. S2), which indicated that *LlWOX9* and *LlWOX11* maintain the normal division of the meristem.

A recent study showed that OsWOX11 can recruit the H3K27me3 demethylase JMJ705 to activate the expression of related genes during rice shoot development (Cheng *et al*., 2018). We speculate that LlWOX11 may regulate the expression of downstream genes through a similar mechanism to promote the formation of bulbils.

### *Cytokinins induce the transcription of* LlWOX9 *and* LlWOX11 *through type-B LlRRs*

Many studies have shown that *WOX* family genes can be induced by plant hormones, such as auxin, cytokinin and gibberellin (Gonzali *et al*., 2005; Leibfried *et al*., 2005; Weijers *et al*., 2006; Sarkar *et al*., 2007; Skylar *et al*., 2010). Cheng *et al*. (2014) analysed the promoters of rice *WOX* family genes and found that there are abundant hormone response elements in these promoters, with the promoter regions of all family members including cytokinin response elements (NGATT/C) and auxin response elements (TGTATC or GAGACA). Further study showed that *OsWOX5*, *OsWOX11*, *OsWOX12A* and *OsWOX12B* could be rapidly induced by NAA and 6-BA (Cheng *et al*., 2014). In our study, we also found a large number of plant hormone response elements, especially cytokinin response elements, in the *LlWOX9* and *LlWOX11* promoters. Then, we demonstrated that the expression of *LlWOX9* and *LlWOX11* could be induced by cytokinin, during which the expression of *LlWOX9* reached a peak after 2.5 h of induction, and the expression of *LlWOX11* reached its highest level after 2 h of induction.

### LlWOX11 *mediates cytokinin signalling by inhibiting the transcription of* LlRR9

Some studies have shown that both *WOX9* and *WOX11* can mediate cytokinin pathways to regulate plant development (Wu *et al*., 2005, 2007; Zhao *et al*., 2009; Wang *et al*., 2014c; Jiang *et al*., 2017). In rice crown root formation, OsWOX11 can directly bind and inhibit the transcription of *OsRR2* to mediate cytokinin signalling (Zhao *et al*., 2009). *OsRR2* is specifically expressed in the crown root and is a member of the type-A RRs, which are negative regulators of cytokinin signalling. Therefore, after the transcription of *OsRR2* is inhibited, cytokinin signalling is enhanced to induce crown root formation (Zhao *et al*., 2009). The ERF3 protein can bind to the OsWOX11 protein and further enhance the transcriptional inhibition of *OsRR2* by OsWOX11 (Zhao *et al*., 2015). We identified a similar mechanism, as our results showed that LlWOX11 can directly bind the promoter of *LlRR9*, a type-A *LlRR* gene, and inhibit its transcription to enhance cytokinin signalling and thus promote bulbil formation.

*WOX9* mediates cytokinin pathway signalling in a different way. During *A. thaliana* seedling development, WOX9 seems to promote the expression of type-A *RR*s. In the *wox9* mutant, *ARR5* expression is decreased, which causes shoot meristem development termination (Skylar *et al*., 2010; Skylar & Wu, 2010). In rice, OsWOX9 plays a negative role in regulating the expression of type-A *RR*s (Wang *et al*., 2014c). A study in rice revealed that *OsWOX9* modulates the cytokinin pathway to regulate the growth height and flowering time of tillers and main branches (Wang *et al*., 2014c). In the *Oswox9* mutant, tiller elongation is inhibited, and in the shortened internodes, the expression of *OsCKX4*, *OsCKX9* and several type-A *OsRR* genes (*OsRR6*, *OsRR9*, *OsRR10*) is increased (Wang *et al*., 2014c). However, in our study, we did not find an effect of LlWOX9 on the expression of *LlRR9*.

### LlWOX9 *may mediate gibberellin signalling*

In the *Oswox9* mutant, the tillers have shorter internodes with cells that are fewer in number and unelongated relative to those of the wild type, and *OsWOX9* activity in internode elongation is directly or indirectly associated with GA signalling (Wang *et al*., 2014c). The organ boundary gene *ARABIDOPSIS THALIANA HOMEOBOX GENE 1* (*ATH1*) and the gibberellin signalling *DELLA* genes maintain the compressed rosette growth habit of *Arabidopsis*. The loss of *ATH1* and *DELLA* function causes a change from a rosette to caulescent growth habit (Ejaz *et al*., 2021). The phenotypes of *LlWOX9*-overexpressing *Arabidopsis* lines show elongated internodes, and we speculate that *LlWOX9* may regulate the internode elongation associated with GA signalling.

### LlWOX9 *and* LlWOX11 *may be involved in scale development and anthocyanin synthesis*

A recent study on the genome of garlic bulb plants showed that two *WOX* family genes (*Asa7G00799.1* and *Asa3G03517.1*) are involved in bulb development, among which *Asa7G00799.1* is expressed specifically in bulbs and positively correlated with bulb weight (Sun *et al*., 2020). Our results showed that *LlWOX9* and *LlWOX11* were highly expressed in scales and that the expression of *LlWOX9* and *LlWOX11* reached the highest level at the stage of bulbil scale development (S5), indicating that *LlWOX9* and *LlWOX11* may be involved in the development of lily bulbs.

Interestingly, we found that the overexpression of *LlWOX9* and *LlWOX11* not only promoted bulbil formation but also resulted in abnormal purple-black bulbils on the leaf axils (Fig. S2). Although bulbils may gradually turn purple-black during development under normal circumstances, the overexpression of *LlWOX9* and *LlWOX11* significantly advanced this change. A recent study showed that *PgWOX11* in *Panax ginseng* can positively regulate the expression of *ERF1B* (an *AP2/ETHYLENE-RESPONSIVE FACTOR*) and thus regulate the biosynthesis of ginsenosides (Liu *et al*., 2020a,b). Therefore, we speculate that *LlWOX9* and *LlWOX11* may be involved in the synthesis of anthocyanins.

## Conclusion

In conclusion, we revealed the molecular mechanism by which WOX genes cooperate with cytokinin signalling to regulate bulbil formation. Type-B LlRRs promote the transcription of *LlWOX9* and *LlWOX11*, and LlWOX11 inhibits the transcription of type-A *LlRR9* to enhance cytokinin signalling, thus promoting bulbil formation (**Fig. 9**).

**Fig. 9.**
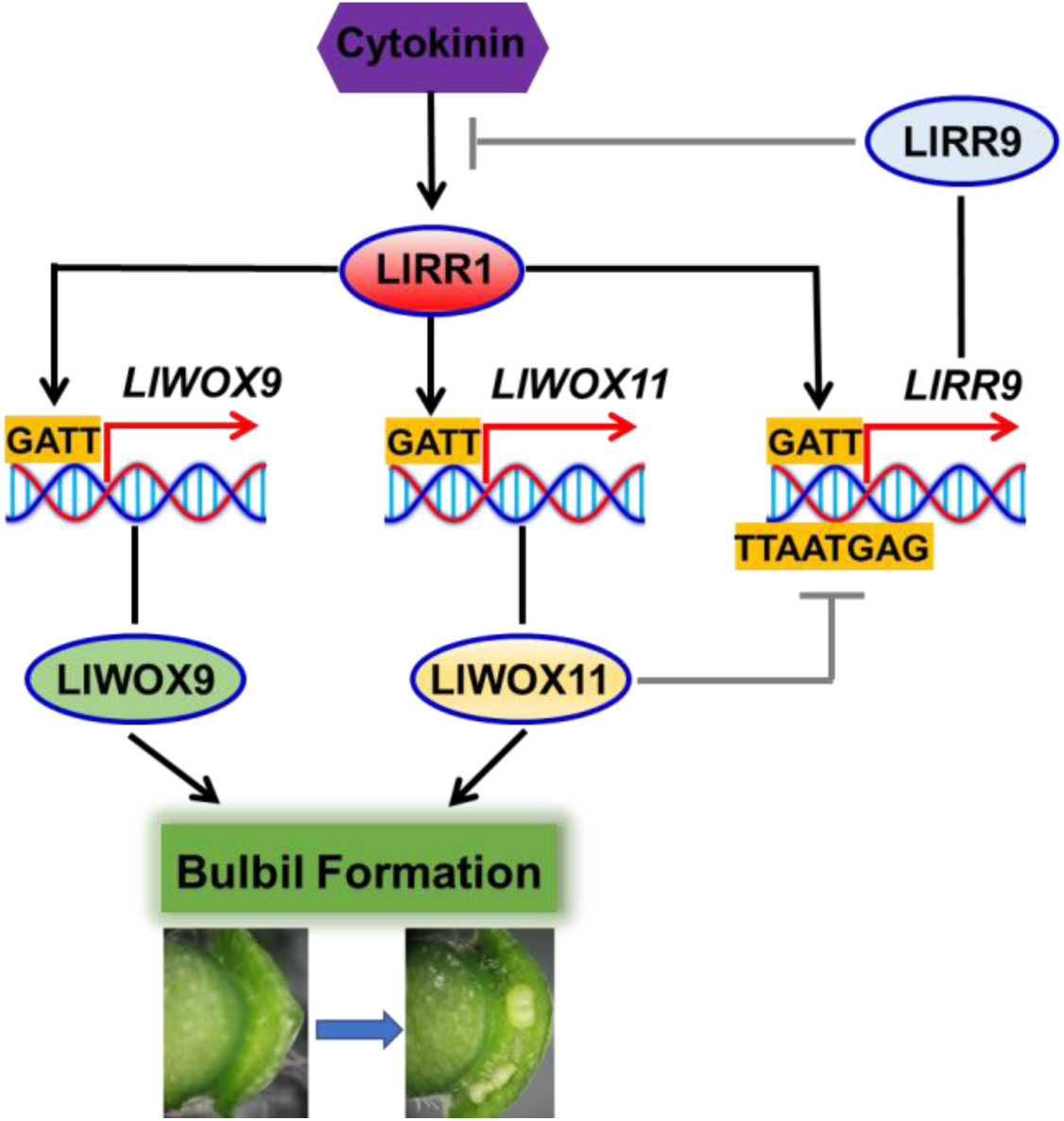
Model of WOX gene cooperation with cytokinin signalling to regulate the bulbil formation.

## Acknowledgements

We acknowledge Xia Cui (Chinese Academy of Agricultural Sciences, China) for the pluc-35Rluc vector and technical assistance. This work were supported by National Natural Science Foundation of China (31902043), Science and technology projects of Guizhou Province (20201Y121), National key R & D program of China (2019YFD1001002) and the Central Public-interest Scientific Institution Basal Research Fund (IVF-BRF2021017).

## Author contribution

JM and PY designed the research. GH, YC, YT, LW, MS, JW, and LX conducted the experiments. GH analyzed the data and wrote the manuscript. All authors read and approved the manuscript.

## Accession numbers

RNA-seq raw reads from this article can be found in the NCBI SRA data under accession number SRP103184. Gene accession numbers used in this study: LlRR1 (MW509629), LlRR2 (MW509630), LlRR10 (MW509631), LlRR11 (MW509632), LlRR12 (MW509633) and LlRR9 (MW509634).

## Supporting information

**Table S1.** Primers used in full length and promoter sequence cloning of *LlWOX9* and *LlWOX11*.

**Table S2.** Primers used in qRT-PCR.

**Table S3.** Primers used in vectors construction.

**Fig. S1.**
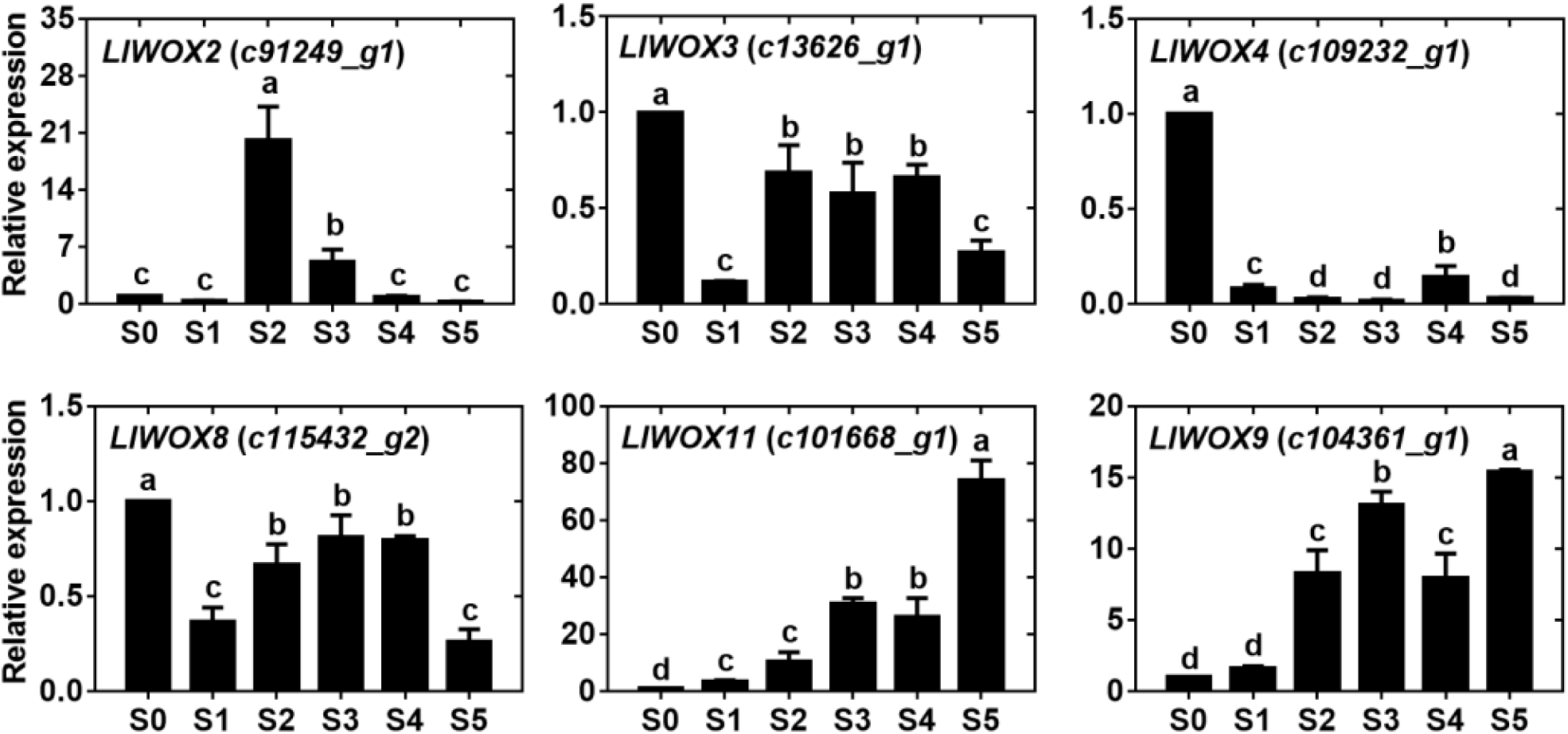
The expression of *WOX* genes during bulbil formation. Values are means ± SDs (n=3). Lowercase letters (a-d) indicate statistically significant differences at *P* < 0.05.

**Fig. S2.**
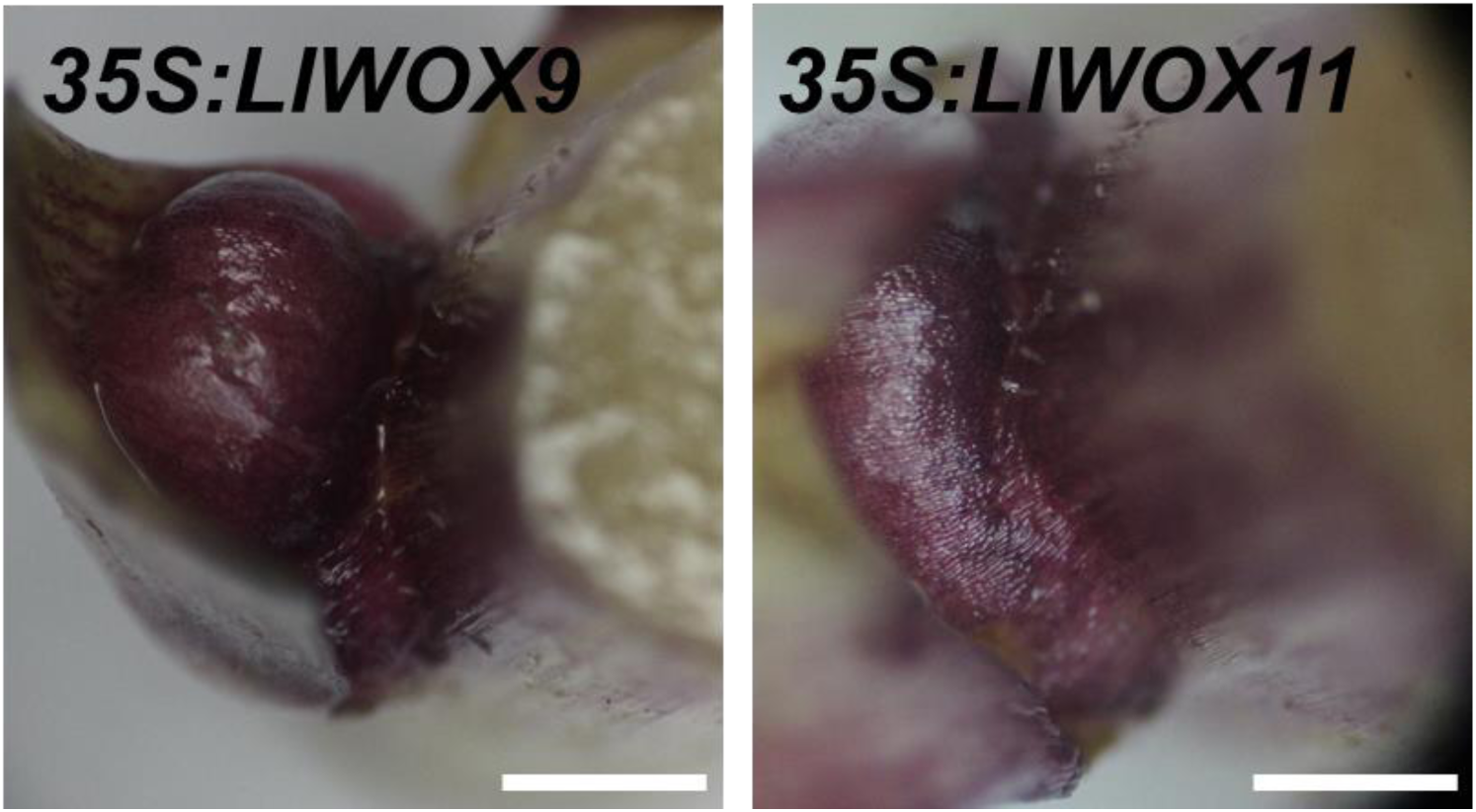
The phenotype of abnormal proliferation in leaf axil after overexpression of *LlWOX9* and *LlWOX11*.

